# Postural adjustments in anticipation of predictable perturbations allow elderly fallers to achieve a balance recovery performance equivalent to elderly non-fallers

**DOI:** 10.1101/506501

**Authors:** Charlotte Le Mouel, Romain Tisserand, Thomas Robert, Romain Brette

## Abstract

**Background:** In numerous laboratory-based perturbation experiments, differences in the balance recovery performance of elderly fallers and non-fallers are moderate or absent. This performance may be affected by the subjects adjusting their initial posture in anticipation of the perturbation.

**Research questions:** Do elderly fallers and non-fallers adjust their posture in anticipation of externally-imposed perturbations in a laboratory setting? How does this impact their balance recovery performance?

**Methods:** 21 elderly non-fallers, 18 age-matched elderly fallers and 11 young adults performed both a forward waist-pull perturbation task and a Choice Stepping Reaction Time (CSRT) task. Whole-body kinematics and ground reaction forces were recorded. For each group, we evaluated the balance recovery performance in the perturbation task, change in initial center of mass (CoM) position between the CSRT and the perturbation task, and the influence of initial CoM position on task performance.

**Results:** The balance recovery performance of elderly fallers was equivalent to elderly non-fallers (p > 0.5 Kolmogorov-Smirnov test). All subject groups anticipated forward perturbations by shifting their CoM backward compared to the CSRT task (young: 2.1% of lower limb length, elderly non-fallers: 2.7%, elderly fallers: 2.2%, Hodges-Lehmann estimator, p < 0.001 Mann-Whitney U). This backward shift increases the probability of resisting the traction without taking a step.

**Significance:** The ability to anticipate perturbations is preserved in elderly fallers and may explain their preserved balance recovery performance in laboratory-based perturbation tasks. Therefore, future fall risk prediction studies should carefully control for this postural strategy, by interleaving perturbations of different directions for example.

## I. Introduction

With aging, there is an increasing incidence of falling (1), causing dramatic impact on health and quality of life (2). As falls may occur even in elderly adults who appear healthy (1), it remains very difficult to predict if and when an elderly person will fall. An observational study of falls in elderly people residing in long-term care identified that around a third of these falls occurred when the person failed to respond appropriately to a perturbation (following a trip, stumble, hot or bump) (3). Therefore, a major focus of research has been to characterize the balance responses of elderly fallers and non-fallers to controlled external perturbations in a laboratory environment. Although certain studies show moderate differences between elderly fallers and non-fallers (4–7) there is a surprising number of prospective studies that fail to show any difference between future fallers and non-fallers (8–11). These studies revealed marked differences between young and elderly subjects, therefore it suggests that the postural responses of elderly fallers and non-fallers to external perturbations might simply not differ under the experimental conditions. Due to the repetition of similar perturbations in controlled laboratory environments, it is possible for the subjects to anticipate essential aspects of the upcoming perturbation (direction, timing, amplitude, etc.). The ability to anticipate provides several advantages for improving the response to external perturbations which may otherwise lead to a loss of balance (12).

Our hypothesis is that elderly fallers can perform as well as elderly non-faller in certain perturbation tasks because they adjust their initial posture to the perturbation direction. Indeed, a recent theory of postural control emphasizes that subjects can use their own weight to resist an external perturbation by shifting their center of mass (CoM) in the opposite direction to the perturbation force, prior to perturbation onset (13). Knowing the direction of the perturbation in advance is therefore a strong advantage that humans can use to improve their balance responses. Indeed, young adults exposed to repeated backward translations of the support surface (equivalent to a forward push applied on the CoM) gradually shift their CoM backward and improve their ability to resist the perturbation without taking a step (14). It is however not known whether and to what extent elderly adults adjust the position of their CoM in anticipation of external perturbations, and how this impacts their balance performance. Although elderly adults are able of adjusting their standing CoM position when explicitly instructed to do so, the range of standing postures which they can adopt is more limited than young adults (15,16). This suggests that the capacity for adjusting CoM position may be reduced in the elderly compared to the young. However, since elderly fallers can perform as well as elderly non-fallers in certain perturbation tasks, we hypothesize that the capacity to adjust their posture before an external perturbation with at least partly known characteristics is preserved in elderly fallers relative to non-fallers, attenuating the differences in balance recovery performance between these two groups.

Young adults, elderly non-fallers and elderly fallers participated in a forward waist-pull perturbation experiment. We quantified the balance recovery performance and determined the initial position of the CoM adopted by the different groups. We determined the effect of backwards leaning on task performance, with the hypothesis that leaning backwards would improve task performance, since this allows the person to use their own weight to resist the perturbation. For comparison with a task where leaning backward would be disadvantageous, we also assessed the initial posture of the same subjects during a Choice Stepping Reaction Time (CSRT) task (15). We then quantified the change in the subjects’ initial posture between the two tasks.

## II. Methods

### 1. Protocol

#### a) Population

Eleven young (aged less than 30) and thirty-nine elderly (aged more than 70) healthy subjects without neurological, musculoskeletal or sensorial disorders participated in the study. Details of the exclusion criteria are provided in the Supplementary Methods (I.1). Elderly subjects completed the Activities Specific Balance Confidence questionnaire (16) to estimate fear of falling. Additionally, they were retrospectively classified as fallers if they had experienced at least one fall in the previous year (17). Characteristics of all subjects are summarized in Table 1. Written informed consent approved by the Comité de Protection des Personnes Lyon Sud Est III (study number 2014-A00179-38) was obtained prior to the experiment.

**Table 1.**
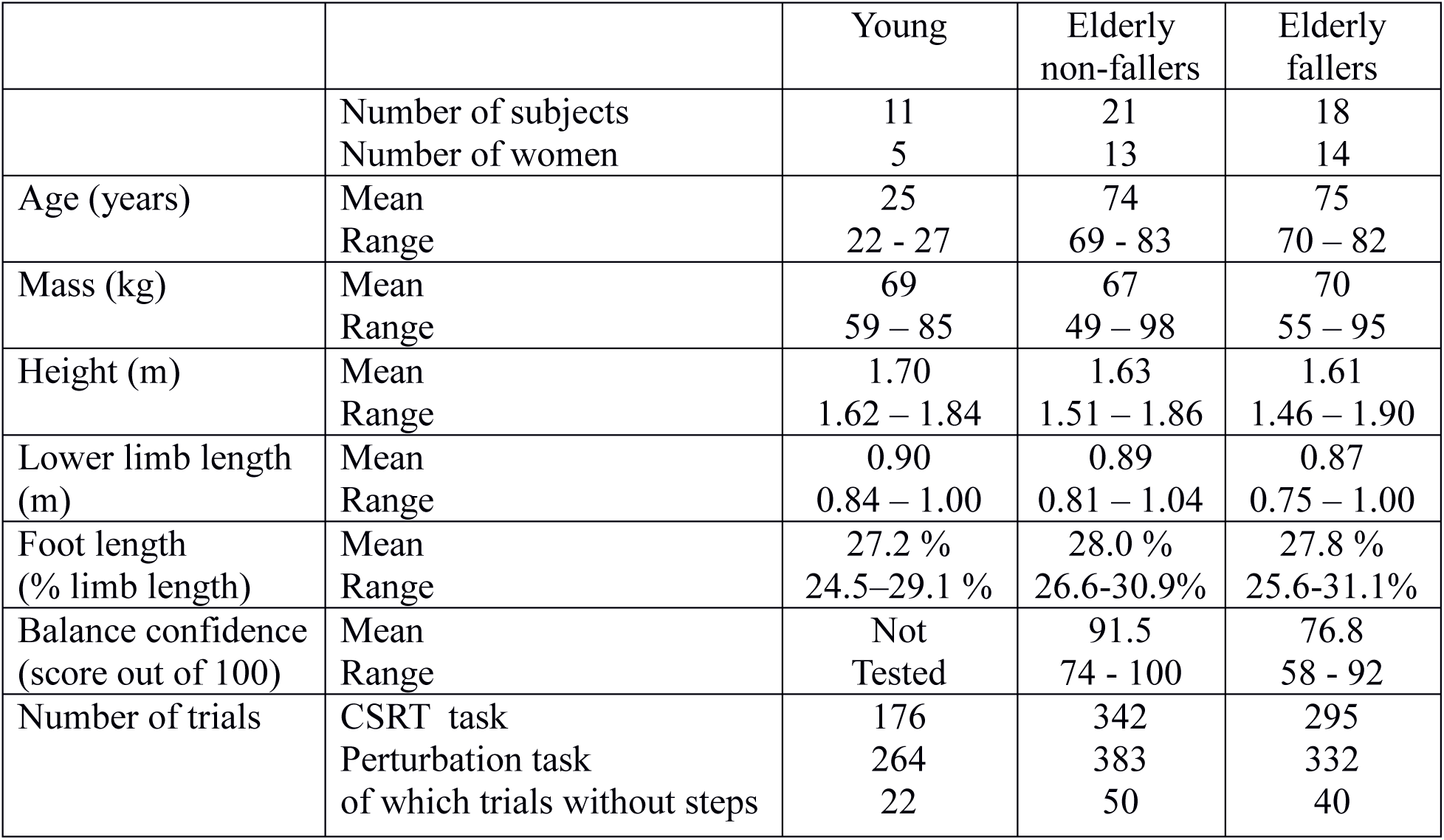
Characteristics of the subjects

#### b) Experimental setup

The lower limb length *L* of each subject was measured as the vertical distance between the great trochanter and the base of the foot. The foot length was measured as the distance between the heel and the longest toe. Subjects were equipped with six reflective markers on each foot, whose positions were recorded using eight cameras (Eagle 1.3 Mpx, Motion Analysis^(R)^, Santa Rosa, CA, USA) sampled at 100 Hz. We used the markers positioned on the tip of the second toe to determine the forwards edge of the foot at trial onset. To estimate the balance performance in perturbation trials, we used the trajectory of the fifth metatarsal joint markers, because they were never masked from the camera after perturbation onset. Four force platforms (60 cm × 40 cm, Bertec^(R)^, OH, USA) were used to record ground reaction forces and torques sampled at 1000 Hz. All subjects performed the CSRT task first, and then the perturbation task. For each trial, subjects initially stood quietly with one foot on each of the two back platforms (Figure 1, A-B).

**Figure 1.**
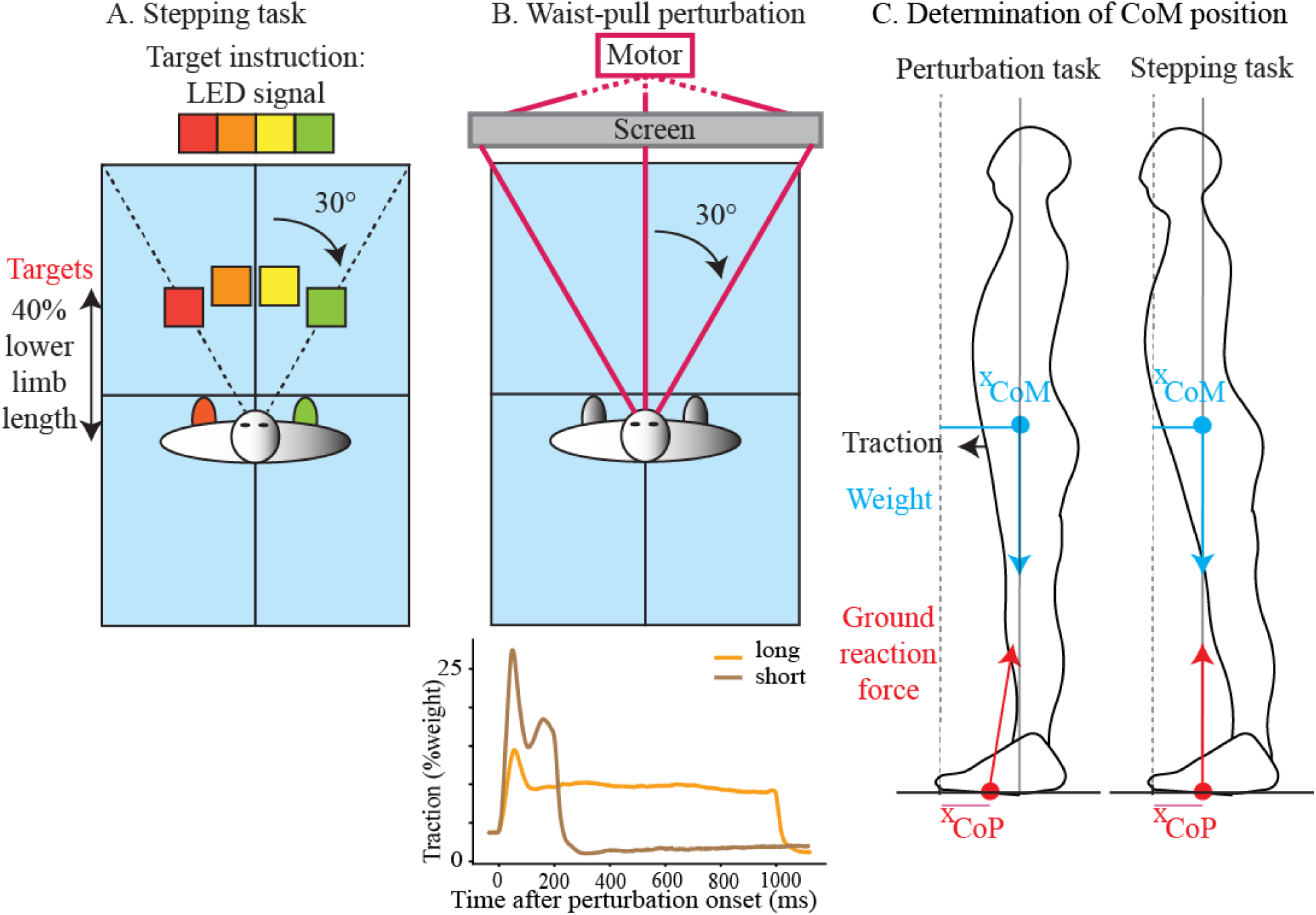
Methods. A.-B Protocol: the subject initially stands with one foot on each of the two back force platforms out of four (blue). A. In the Choice Stepping Reaction Time task, four stepping targets are placed in front of the subject at a distance of 40% of the subject’s lower limb length. Two of these targets are located centrally (yellow and orange), and the two others are located laterally, at an angle of 30° (red and green). Four light-emitting diodes (LED), corresponding to the four targets, are placed in front of the subject to indicate on which target the subject should step. B. (Upper) In the perturbation task, three cables are attached to the subject’s harness at waist level and, on any given trial, only one of these cables is attached to a motor hidden behind a screen. (Lower). The perturbation is proportional to the subject’s weight and is either short (brown) or long (yellow). C. The person is assumed to be in a quasi-static posture before trial onset. For the perturbation task (left), due to the initial pre-tension (black arrow), the CoM (blue dot) is backward of the CoP (red dot). For the CSRT task (right), the CoM and CoP are aligned.

The protocol and results of the CSRT are published elsewhere (15). Briefly, four stepping targets and four corresponding light-emitting diodes (LED) were placed in front of the subject (Figure 1.A). Subjects were instructed to: “place [their] corresponding foot as fast as possible onto the corresponding target indicated by the LED.” Leftwards targets (red and orange) were to be stepped on with the left foot, and rightwards targets (yellow and green) with the right foot. Each subject performed one block of 16 trials, with each target presented four times, in random order. Trials in which the subject appeared to hesitate for a long while, or in which the subject stepped with the wrong foot, were repeated at the end of the initial sequence of 16 trials.

In the perturbation task, subjects wore a safety harness to prevent injury in case of a fall. Three cables were attached to the harness at waist level. Each cable could pull the person either straight forward or laterally at an angle of 30° (Figure 1.B, upper panel). On any given trial, one of the three cables was attached to a rotating motor which, after a random waiting period between 1 and 12 seconds, applied a force controlled in amplitude and duration. The applied force was either “short” (200 ms duration with a peak amplitude of 27% of the subject’s weight, brown curve in Figure 1.B, lower panel), or “long” (1000 ms duration with a peak amplitude of 14% of the subject’s weight, orange curve). The integral of the force applied during the long perturbation was 91% larger than the integral of the short perturbation. To avoid any whipping effect when stretching the cable, an initial pretension was applied by the rotating motor before perturbation onset. The two other cables were stretched by small mass equivalent to the pre-tension force attached to their end, resulting in a total pretension of 3.5% of the subject’s body weight. Additional details can be found in the Supplementary Methods (I.2). Subjects were instructed to “recover balance as fast as possible and in the shortest possible distance”. Elderly subjects performed 18 trials, with each of the six perturbations (three directions, two amplitudes) presented three times, in random order. Young subjects performed 24 trials, with each perturbation presented four times, in random order.

### 2. Analysis

#### a) Performance in perturbation trials

Performance was assessed by the distance between the initial foot position and the final foot position, measured once a steady balance was recovered. The shorter the distance, the better the task success. For comparison across subjects, this distance was normalized to the subject’s lower limb length. The position of the fifth metatarsal joint of each foot was used to indicate foot position (error of measurement < 1 mm, i.e. 0.1% limb length). This method is illustrated in the Supplementary Methods (II.1). The distribution of performance distances was bimodal (Figure 2.A-D), and in trials for which this distance was inferior to 8% of the person’s lower limb length, it was considered that the subject did not step (non-step trial). We then analyzed two variables: the frequency of non-step trials, and the performance distance in which the subject took at least one step (step trials).

**Figure 2.**
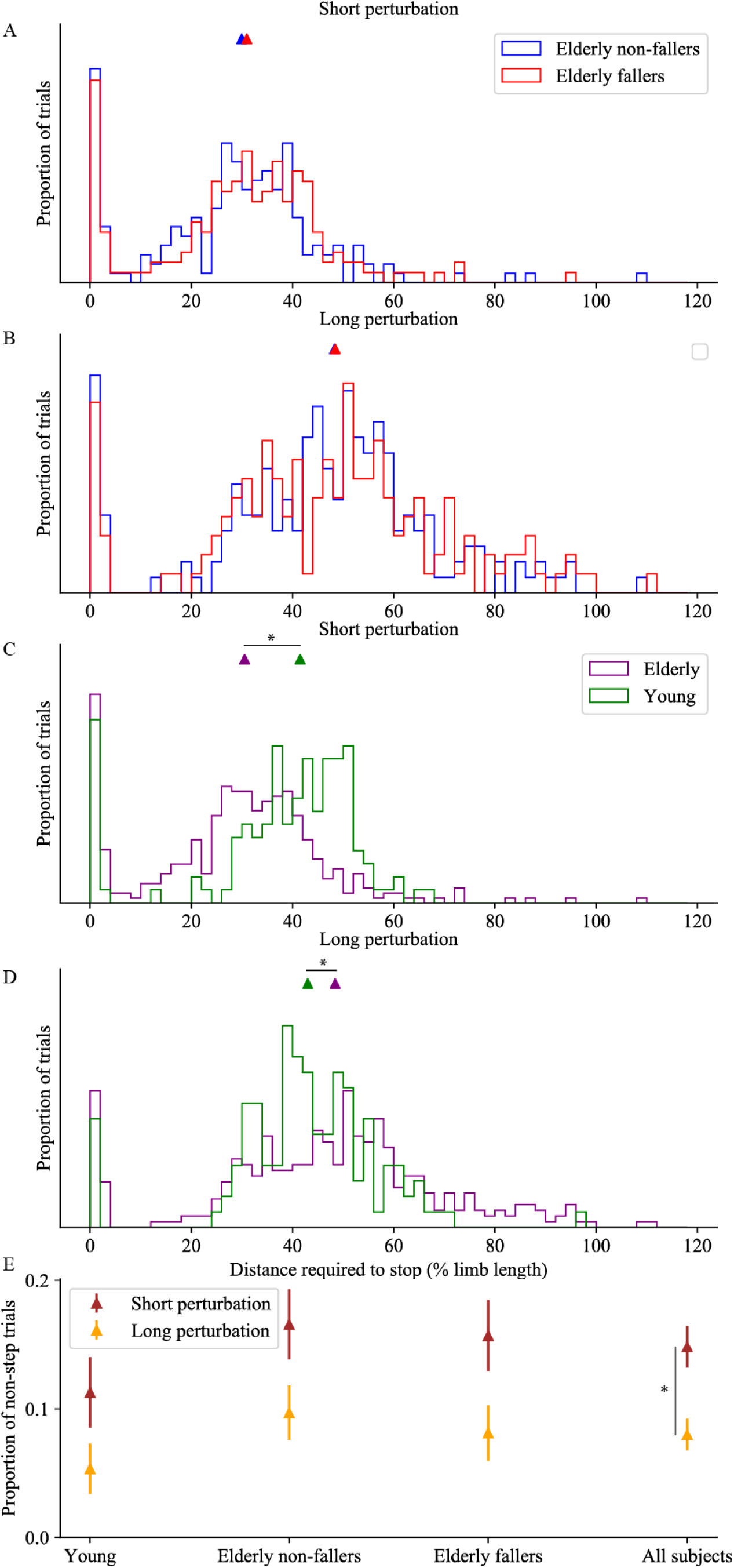
Performance in the perturbation task. Graphs show the normalized distribution of distance required to stop the motion triggered by the perturbation, normalized to the subject’s lower limb length, for short (A, C) and long (B, D) perturbations. In panels A. and B., elderly non-fallers are shown in blue and elderly fallers in red. In panels C. and D., elderly fallers and non-fallers are pooled (purple), for comparison with the young subjects (green). For each group and perturbation duration, the median distance is indicated as a triangle on the top of the graph. E. Proportion of non-step trials for short (brown) and long (orange) perturbations; error bars indicate the standard error on the estimation of this proportion (additional details are provided in the Supplementary Methods I.3). Statistically different results are indicated as a star.

#### b) Initial posture preceding stimulus onset

We determined the subject’s posture in the 500 ms preceding the onset of the LED in the CSRT task, and in the 500 ms preceding the onset of the perturbation in the perturbation task. During this waiting period, the subject is in quasi-static standing posture, therefore the torques acting upon the body cancel each other out. The horizontal distance between the CoM and the center of pressure (CoP) is determined by the torque of the external forces acting on the body other than weight and the ground reaction force (13). The position of these points within the feet remains however free, allowing subjects to adopt different initial postures. We determined the front of the feet as the mid-point between the markers on the toes of the left and right feet. We then used the forceplates to determine the antero-posterior positions of the CoP and CoM relative to the front of the feet, *x*_*CoP*_ and *x*_*CoM*_.

In the perturbation task, because an initial pretension was applied by the cables to the subject in the forward direction, the CoM was positioned backward of the CoP (Figure 1.C, left panel). This pretension was assumed to be applied horizontally, at a height equal to the subject’s lower limb length *L*. Its amplitude was assumed to be on average opposite to the mean anteroposterior ground reaction force (*F*_*x*_) measured by the forceplates. The sum of torques is then:

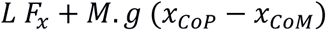

Where *M.g* is the person’s weight, defined as the mean vertical ground reaction force during the waiting periods in the CSRT task. We assumed that the sum of torques was on average null during each waiting period to determine *x*_*CoM*_ from *x*_*CoP*_ and *F*_*x*_. To adequately compare the initial postures measured in the CSRT and perturbation tasks, the same method was used to determine *x*_*CoM*_ in the CSRT task.

The perturbation amplitude was scaled to the subject’s weight, therefore its torque was proportional to *L M g*. To compare torques across subjects, we normalized *x*_*CoM*_ by the subject’s lower limb length, rather than by the subject’s foot length as is commonly done (18,19). This method is illustrated in the Supplementary Methods (II.1).

#### c) Error of measurement

The error of measurement of *x*_*CoM*_/*L* is the sum of the error of measurement of the positions of the toe markers, and of the error due to the force platform measurement. The former has a constant bias due to the unreliable placement of the markers on the anatomical landmarks, and a variable noise component due to the noisy measurement of the marker positions. Overall, the error of measurement of the motion capture device is inferior to 1 mm, i.e. 0.1% limb length. The latter may have a constant bias due to the calibration of the force platforms, and has a variable noise component due to slow changes in posture which were estimated to be inferior to 0.2% of lower limb length (see Supplementary Methods II.2). When considering differences in *x*_*CoM*_/*L* between two trials of a given subject, the constant biases cancel out. The error of measurement for differences in posture is therefore only affected by the variable noise components, estimated to be inferior to 0.3% of lower limb length per trial. To report differences in posture between two samples of size *n*_1_ and *n*_2_, we indicated the standard error of the mean as 0.3% 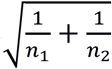

The error in measurement of the distance required to recover balance is only due to the noise in the marker positions (inferior to 0.1% limb length). When reporting differences in balance performance, we used 0.1% 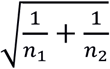

### 3. Statistical analysis

The 2-proportion z-test was used to compare the proportion of non-step trials across subject groups and across perturbations.

The Mann-Whitney U-test was used to compare differences between the medians of continuous distributions. It was first used to compare the median performance in the perturbation task across subject groups and across perturbations. It was then used to compare the median initial *x*_*CoM*_ between the CSRT task and the perturbation task, within each subject group. Finally, it was used to compare the median initial *x*_*CoM*_ for perturbation trials with and without steps, within each subject group.

When a statistical difference between two medians was found, this difference was estimated using the Hodges-Lehmann estimator.

When no statistical difference between two medians was found, as for instance in the balance performance of elderly fallers and non-fallers in the perturbation task for each perturbation amplitudes, then a Kolmogorov-Smirnof test was used to determine whether there was any difference in the distributions.

For each elderly subject, the Hodges-Lehmann estimator was used to estimate the subject’s change in *x*_*CoM*_ between the CSRT and the perturbation task. A linear regression was used to determine whether this change was correlated to fear of falling within fallers, within non-fallers, and across elderly subjects.

Additional details can be found in the Supplementary Methods (II.3).

## III. Results

### 1. Balance performance in the perturbation task

The distribution of the distances required to recover balance is illustrated in Figure 2.A-D for each group and the two perturbations. In trials with at least one step, this distance was 33% of lower limb length for elderly non-fallers and 34% for elderly fallers (short perturbation) and 50% for both elderly non-fallers and fallers (long perturbation). We found no significant difference between the performance of the elderly fallers (red) and elderly non-fallers (blue), regardless of the perturbation (p > 0.5 for both short and long perturbations).

Elderly fallers and non-fallers were pooled together (purple trace) and compared to young subjects (green). For young subjects, the distance required to stop was 43% of lower limb length for both short and long perturbations. For short perturbations (Figure 2.C), this distance was significantly larger than for elderly subjects (p < 0.001) by 9.3% ± 0.01% of lower limb length. For long perturbations (Figure 2.D), this distance was significantly smaller than for elderly subjects (p < 0.001) by 5.4% ± 0.01% of lower limb length.

Across all subjects, the proportion of non-step trials was significantly larger for short compared to long perturbations (p < 0.001, see Figure 2.E). Within each perturbation type, the proportion of non-step trial was slightly smaller for young compared to elderly but this difference was not significant. No difference was observed between elderly fallers and elderly non-fallers (p > 0.1).

### 2. Change in posture across tasks

The distribution of distances between the initial CoM position and the forward edge of the feet for each group and both tasks is presented in Figure 3. For the perturbation task, the median initial distance across all subjects was on average 17.5% of the subject’s lower limb length, with no significant differences between groups (p > 0.1). For the CRST task, the median distance was 15.4% of the person’s lower limb length. This distance was slightly smaller in non-fallers (15.2%), compared to fallers (16.0%, p < 0.01) and young adults (15.7%, p < 0.05). No significant difference was found between young adults and elderly fallers (p > 0.5).

**Figure 3.**
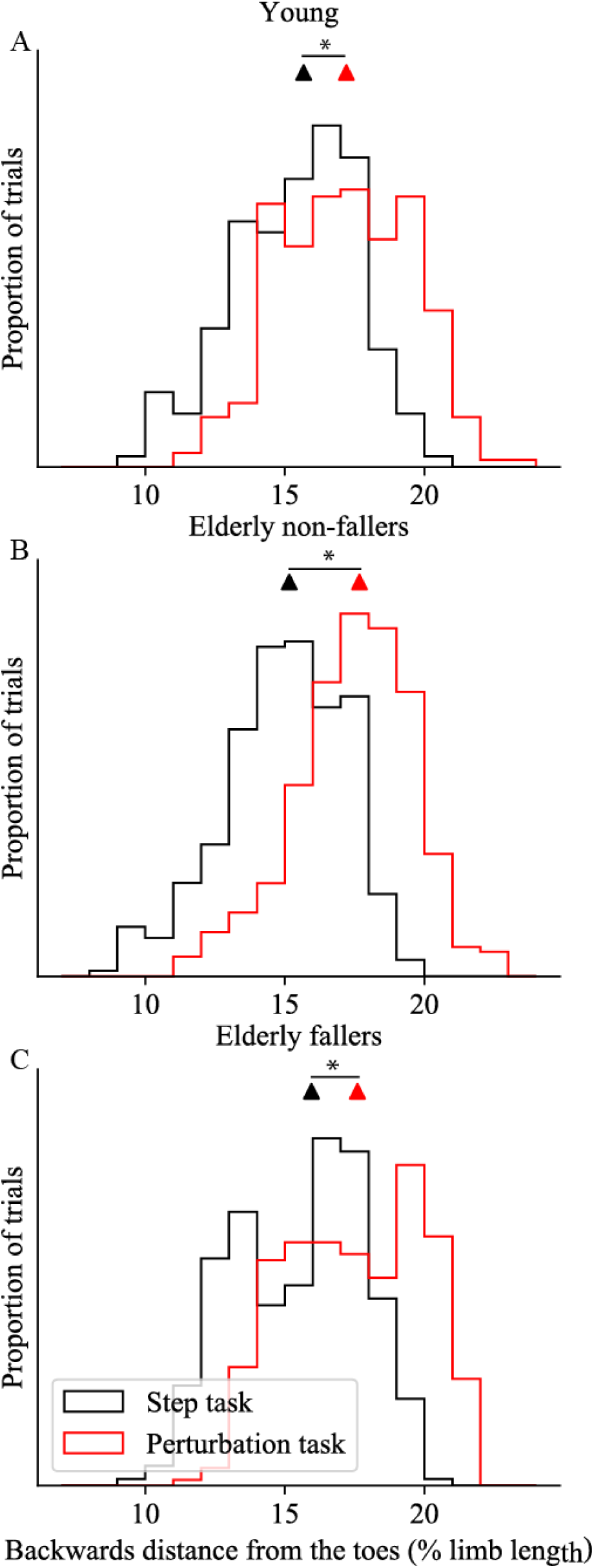
Change in posture across tasks. Graphs show the normalized distribution of the initial position of the CoM backward of the toes as % of lower limb length, in the perturbation task (red) and in the CSRT task (black), for young subjects (A), elderly non-fallers (B) and elderly fallers (C). For each group and task, the median CoM position is indicated as a triangle on the top of the graph. Statistically different results are indicated as a star.

All groups of subjects shifted their CoM backward in the perturbation task compared to the CSRT task (young: p < 0.001, non-fallers: p < 0.001, fallers: p < 0.001). This backward shift is of 1.8% ± 0.03% of lower limb length for the young adults (Figure 3.A), 2.5% ± 0.02% for the elderly non-fallers (Figure 3.B) and 2.0% ± 0.02% for the elderly fallers (Figure 3.C). To provide values comparable with previous studies, we also reported the results in % foot length (Table 2).

**Table 2.** Difference in initial CoM position relative to the toes

|  | In the perturbation task relative to the CSRT task |  | In perturbation trials with steps relative to trials without steps |  |
| --- | --- | --- | --- | --- |
|  | % limb length | % foot length | % limb length | % foot length |
| Young | 1.8 (p < 0.001) | 6.7 (p < 0.001) | 2.8 (p < 0.001) | 7.2 (p < 0.001) |
| Elderly non-fallers | 2.5 (p < 0.001) | 8.9 (p < 0.001) | 0.7 (p = 0.01) | 3.0 (p < 0.001) |
| Elderly fallers | 2.0 (p < 0.001) | 7.6 (p < 0.001) | 2.3 (p < 0.001) | 10.9 (p < 0.001) |

Fear of falling was significantly higher (i.e. a smaller Balance confidence score) in the elderly fallers compared to elderly non-fallers (p < 0.001). However, at an individual level, the change in posture across tasks was not correlated with fear of falling (elderly fallers: p > 0.3, elderly non-fallers: p > 0.7).

### 3. Influence of initial posture on balance performance in the perturbation task

To quantify the effect of the initial posture on balance performance, the initial CoM positions in trials with and without steps were compared (Figure 4). Across trials, the initial CoM positions ranged from 11.2 to 23.6 % of lower limb length (37.9 to 82.6 % of foot length), whereas in trials without steps, subjects leaned backwards by at least 15 % of lower limb length (56 % of foot length). All groups of subjects had a more backward initial CoM position in non-step trials compared to trials with at least one step (young adults: p < 0.01, Figure 4.A; elderly non-fallers: p = 0.01, Figure 4.B; elderly fallers: p < 0.01, Figure 4.C). This difference is of 2.8% ± 0.07% of lower limb length for the young adults, 0.7% ± 0.05% for the elderly non-fallers and 2.3% ± 0.05% for the elderly fallers. Results in % of foot length are indicated in Table 2.

**Figure 4.**
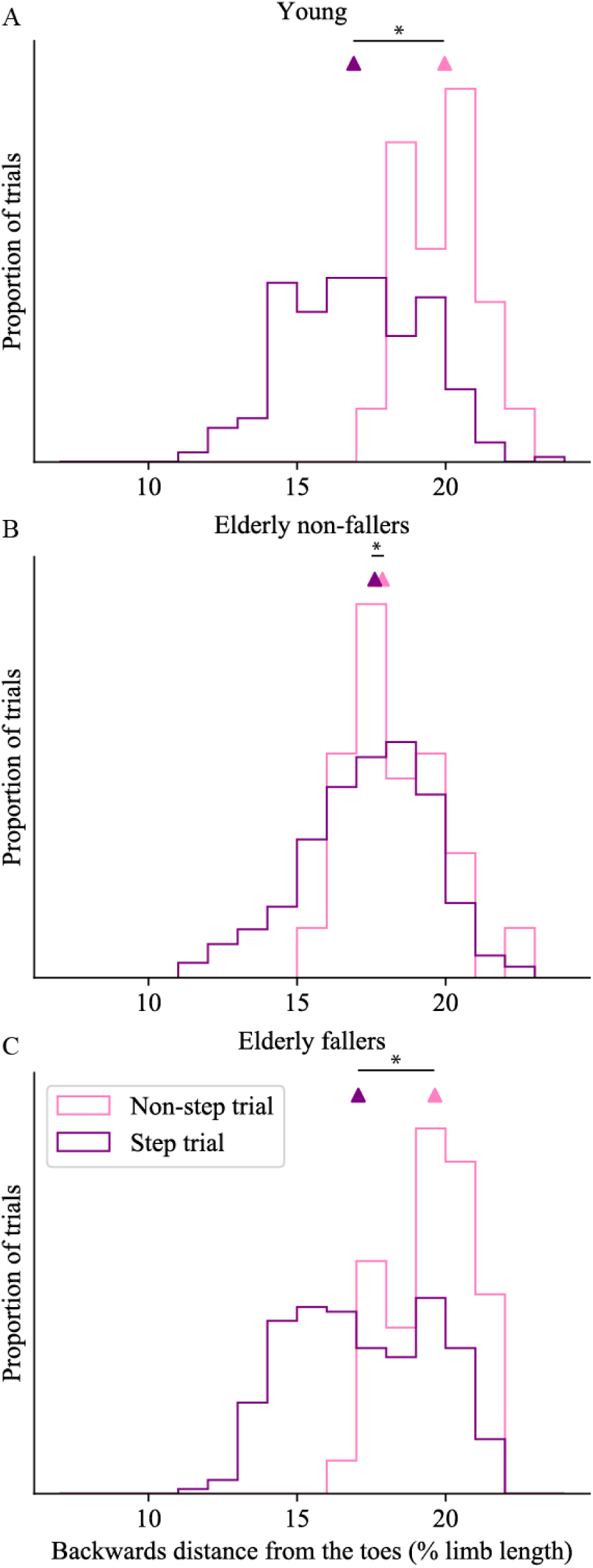
Influence of posture on balance performance. Graphs show the normalized distribution of the initial CoM position in the perturbation task for trials in which the subjects took a step (purple) and for trials in which the subjects did not step (pink), for young subjects (A), elderly non-fallers (B) and elderly fallers (C). For each group and condition, the median CoM position is indicated as a triangle on top of the graph. Statistically different results are indicated as a star.

## IV. Discussion

When exposed to repeated waist-pull perturbations with a predictable dominant forward component, elderly fallers performed as well as elderly non-fallers in terms of distance required to recover balance. This occurred although the perturbation timing, amplitude, duration and lateral direction were not predictable. The absence of difference between the fallers and non-fallers is unlikely to be due only to an inappropriate classification based on a single fall event, because our group of elderly fallers displayed higher fear of falling (Table 1) and reduced performance in the CSRT task (15). Therefore, our study completes the list of studies which have failed to reveal differences in the balance responses between elderly fallers and non-fallers to externally-triggered perturbations in a laboratory setting (8–11).

As in previous laboratory-based perturbation experiments (3), our study revealed differences in balance performance between young and elderly subjects. Young subjects required the same distance to recover balance for both short and long perturbations, although the summed torque induced by the long perturbation was almost twice as large as for the short perturbation. Elderly subjects on the other hand had a more graded response to the perturbation momentum, and required a longer distance to recover balance following the long perturbation than following the short. This suggests that young subjects always trigger the same step (7), regardless of the perturbation momentum. This strategy may be generally effective to recover balance, however for short perturbations it seems to overestimate the necessary step length, as elderly subjects are able to recover balance with shorter distances than young subjects.

In the perturbation protocol used in this study, subjects must produce a backward torque to compensate for the forward torque induced by the pulling cable. This aspect probably appeared very clearly to each subject as soon as the three cables were attached to the harness. The biomechanical action required to produce a backward torque is a shift of the CoP in front of the CoM. However, the CoP cannot be brought further forward than the toes. Therefore, if the person’s feet remain stationary, the maximal backward torque which they can exert is proportional to the distance between their initial CoM position and their toes. Shifting the CoM backward before the onset of the perturbation increases this distance, thereby increasing the potential backward torque needed to resist the perturbation after its onset. This biomechanical requirement explains why we observed a more backward position of the CoM in all subject groups in the perturbation task relative to the CSRT task (Figure 3).

Relative to the natural CoM position adopted in quiet standing, young adults can shift their CoM backwards by up to 20 % of foot length, whereas this distance is reduced to 15 % of foot length in elderly subjects (18,19). In the perturbation task of our study, young subjects shifted their CoM backwards prior to the perturbation onset by 6.7% of their foot length, elderly non-fallers by 8.9 %, and fallers by 7.6%, (Table 2). For elderly subjects this postural shift corresponds to half of their available range. Interestingly, this backwards shift was not correlated to fear of falling.

Shifting the CoM backward prior to an external perturbation is important for resisting the traction (13) and increases the chances to recover balance without taking a step. In our experiment, subjects were instructed to recover balance in the shortest possible distance. Therefore, recovering balance without taking a step was the best possible response. Our results show that initially positioning the CoM further backward increases the proportion of non-stepping trials in all groups of subjects (Figure 4). Trials without steps only occurred when the initial CoM position relative to the toes was larger than 56% of foot length. In quiet standing, young subjects can adopt quiet standing postures with a CoM position relative to the toes ranging from 16.8 % to 76.8 % of foot length, whereas for elderly subjects quiet standing postures range from 29.5 % to 69.5 % (18). The possibility of resisting the traction without taking a step thus requires that subjects initially stand with their CoM relatively close to the backwards edge of their base of support.

Our study demonstrates an anticipatory shift of CoM position preceding perturbation onset when the perturbation direction can be anticipated. This shift was performed by both elderly fallers and non-fallers and helped them to recover balance equally well. Previous studies have demonstrated that anticipatory postural adjustments occur at the initiation of a voluntary movement, such as walking (20) or stepping forwards (15). These adjustments create the necessary momentum to propel the CoM in the direction of the intended movement. For a forwards movement, this is achieved through a backward shift of the CoP after the cue for movement initiation. This CoP shift is of smaller amplitude in the elderly relative to the young (20), and in elderly fallers relative to non-fallers (15), resulting in a slower CoM movement. We thus suggest the ability for slow CoM shifts is preserved in elderly fallers and that what is crucially affected is their ability to rapidly shift the CoM to compensate quickly for a perturbation. Thus, in our perturbation task, subjects are given as much time as they want to prepare for the upcoming perturbation. This task therefore does not probe how fast the subjects can shift their posture, and reveals no difference in performance between fallers and non-fallers. In the CSRT task on the contrary, the subjects do not know with which foot they should step. When the target lights up, they must first shift their weight forward and also onto the stance foot before raising the swing foot. This task probes how fast the subjects can shift their CoM, and elderly fallers perform this task more slowly than elderly non-fallers (15,21). Finally, “incorrect weight shifting”, accounts for 41% of the falls occurring in a nursing home residence (3). Thus, the ability to perform rapid and accurate shifts of their own CoM position appears to be critical in order to identify a potential risk of falling.

According to our results, future studies aiming to predict fall risk in the elderly should be designed to probe the response of the subject to perturbations of unexpected characteristics, including direction. Forward and backward (or leftward and rightward) perturbations should therefore be interleaved in a random manner, to prevent subjects from anticipating on perturbation direction. Thus, a prospective study using waist-pull perturbations in 12 directions revealed differences between future fallers and non-fallers (7). This may be closer to ecological situations where an elderly person encounters an unexpected perturbation requiring a fast postural shift, thus providing a better assessment of fall risk.

