## Supplementary Methods for "Postural adjustments in anticipation of predictable perturbations allow elderly fallers to achieve a balance recovery performance equivalent to elderly non-fallers"

### I. Protocol

#### 1. Exclusion criteria

Young subjects aged less than 30 years were recruited, and were excluded if they had a medical history of musculo-skeletal or neurological problems, or if they were taking medication that could affect motor performance.

Elderly subjects aged at least 70 years were recruited, and were excluded following a medical visit if they met the following criteria:

- Taking more than 3 different psychotropic medications per day
- Regularly using walking aids, such as a crutch or a walking stick
- Severe dyspnea for light efforts
- Congestive cardiac failure at level 3 or more, according to the New York Heart Association standards
- Taking medication for Parkinson or Alzheimer's disease
- A Mini Mental State Examination score of less than 23
- Having experienced a stroke, either in the previous six months, or that left lasting damage
- Having undergone lower limb surgery in the previous six months
- Having visual acuity of less than 2 on the Monoyer scale
- Having a frailty score of 4 or more, according to Fried and colleagues (1).

#### 2. Control of pretension in perturbation trials

In perturbation trials, an initial pretension was applied by the rotating motor so as to avoid a whipping effect when stretching the cable at the onset of the perturbation. To prevent the perturbation direction from being sensed through this initial motor pretension, equivalent masses were attached to the two other cables. This resulted in a total forwards pretension force of around 3.5% of the subject's weight. Thus, on every trial, an experimenter would attach one of the three cables to the motor, and the other two cables to small masses. This was done behind a screen, such that the perturbation direction was not predictable for the subject.

This initial pretension in the cables exerts forward torque on the subject, which must be compensated by a forward position of the CoP relative to the CoM (main text Figure 1.C left panel). The distribution of CoP (Figure 1.A-C) and CoM (Figure 1.D-F) in perturbation trials (red) relative to CSRT trials (black) is shown in Figure 1. Relative to the CSRT trials, subjects compensate for the pretension through a slight forward shift in CoP (Y: 0.4% limb length, $p = 0.1$, O : 0.7%, $p < 0.001$, F : 0.9%, $p < 0.001$), and a larger backward shift in CoM (Y: 1.8% limb length, $p < 0.001$, O : 2.5%, $p < 0.001$, F : 2.0%, $p < 0.001$).

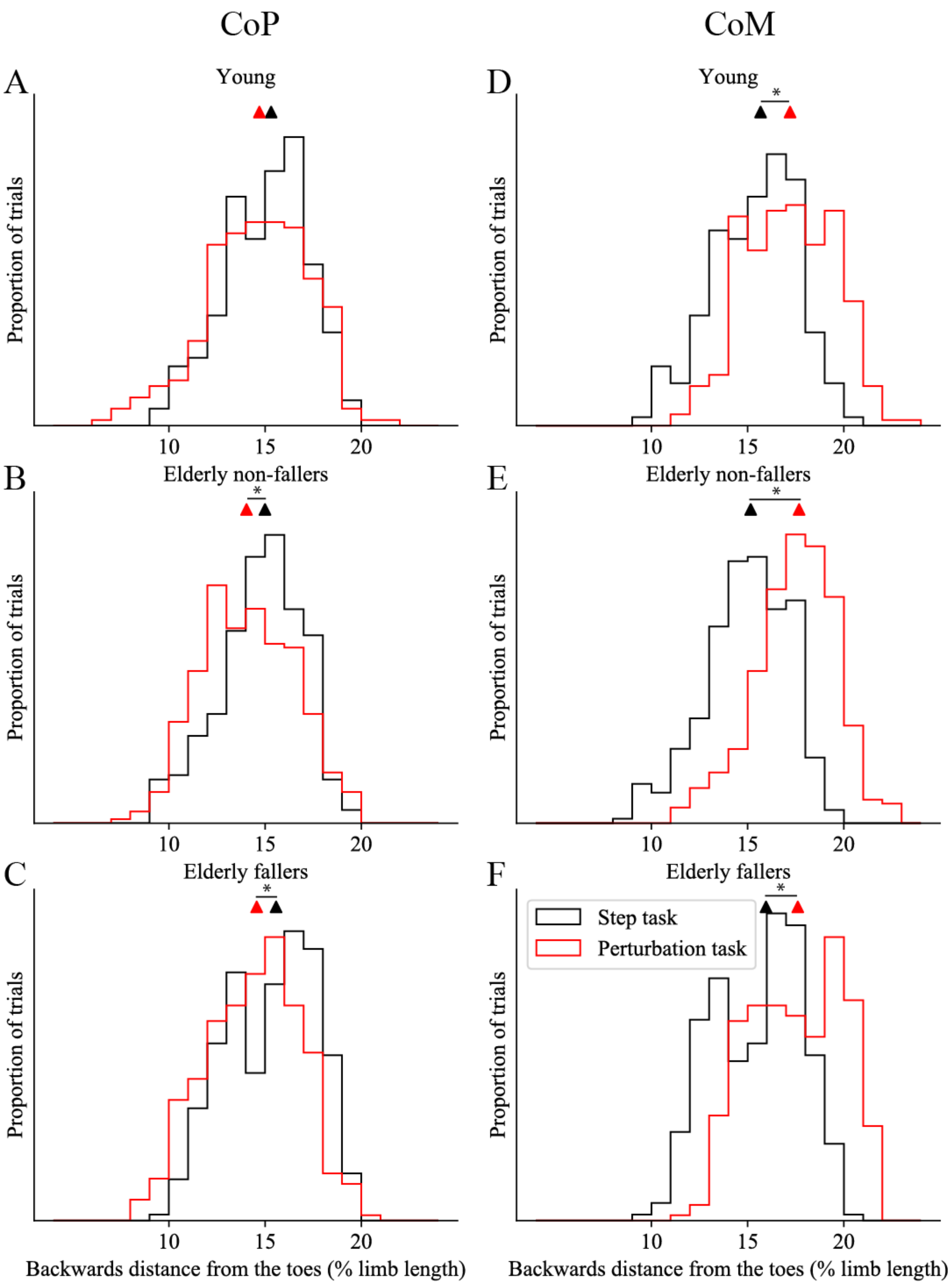


*Figure 1 Initial backward position relative to the toes (normalised by limb length) of the CoP (left column) and CoM (right column) in CSRT trials (black) and in perturbation trials (red), for young (A,D), elderly non-fallers (B, E) and fallers (C, F).*

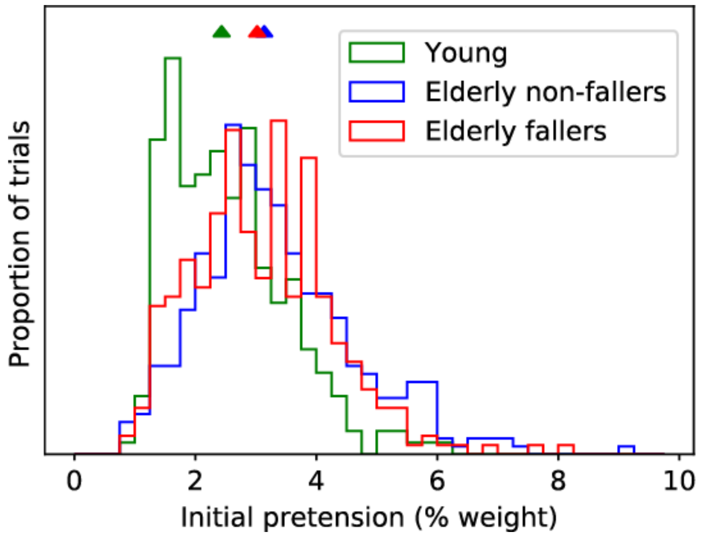

*Figure 2 Initial pretension in perturbation trials (normalized by subject weight) for Y (green), NF (blue) and F (red). Medians of the distributions are shown as triangles.*

The actual value of the pretension was not tightly controlled due to the limitations of the motor applying the traction (which was a step motor), and the distribution of pretensions is shown in Figure 2 (estimated using the average value of the forward component of the ground reaction force in the 500 ms before trial onset). There is no difference in the median pretension between NF and F ($p > 0.4$, Mann-Whitney U). The median pretension is slightly smaller for Y compared to the elderly ($p = 0.03$, Mann-Whitney U, difference of 0.3% subject weight, Hodges-Lehmann estimator).

At a trial by trial level, variations in pretension were not correlated with initial CoM position ($p = 0.06$, $r^2 = 0.004$, linear regression) but were correlated with initial CoP position ($p < 0.01$, $r^2 = 0.18$, linear regression). This suggests that subjects adapt to the perturbation task by shifting their CoM backward by a given amount, then compensate for the variable pretension through a forward shift in CoP relative to the CoM of variable amplitude.

#### II. Analysis

##### 1. Performance in the perturbation task

Subjects were instructed to recover balance as fast as possible and in the shortest possible distance. In the perturbation task, there were a certain number of trials for which the person did not take a step. Given the task instruction, which was to recover balance as fast as possible and in the shortest possible distance, trials without steps should be considered a success. In such trials however, the time at which balance was recovered is ambiguous. Therefore, performance was not determined according to the time required to initiate or perform a protective step, which are commonly used measures of performance in perturbation tasks (2).

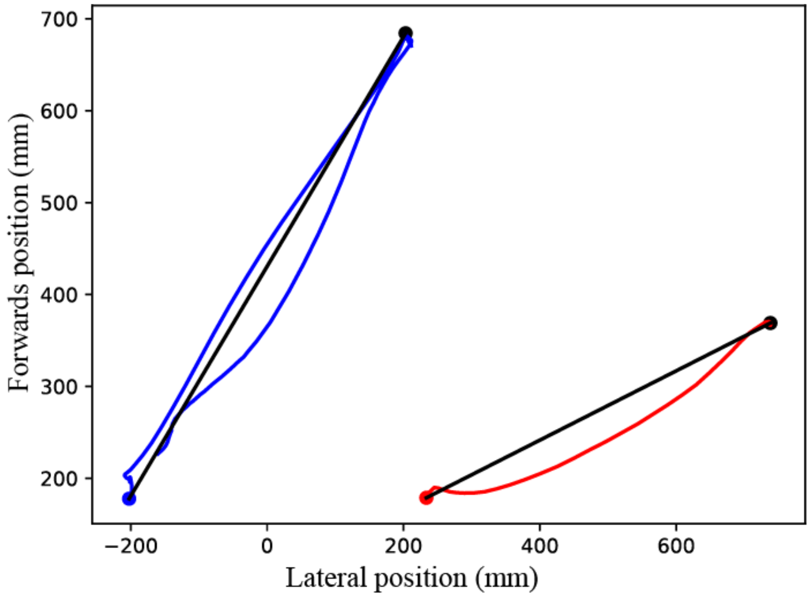


*Figure 3 Calculation of the distance required to stop. The initial position of the left and right feet are shown as a blue and a red dot. The trajectory of the foot after the onset of the perturbation is projected onto the horizontal plane (blue and red curves). The point of the trajectory furthest from the initial position is shown as a black dot.*

Instead, performance was determined as the distance required to recover balance, based on the trajectory of the person's feet. Thus, if the subject's feet did not move, this distance was considered null. If the person took a step, then this distance was the length of the step. In the example presented in Figure 3, the person took two steps forward (one with each foot), then took a step backward with the left foot (blue). The distance required to stop is then considered to be the distance required to stop for the foot which stepped the furthest away from its initial position (here the left foot).

The position of each foot is determined by a kinematic marker placed on the fifth metatarsal joint of the foot. For each foot, the average initial position during the 500 ms preceding the perturbation onset is determined (Figure 3, blue and red dots for the left and right feet). The trajectory of the foot in the 2.5 seconds following the perturbation onset is projected onto the horizontal plane (Figure 3, blue and red curves). The point on this trajectory which is furthest from the initial position is determined (Figure 3, black dots and lines), and this is considered the distance required by that foot to stop. The distance required by the person to stop is then the maximum of the distance required by each foot. Finally, this distance is normalized to the subject's lower limb length for comparison across subjects.

2. Error of measurement of the CoM position

a) High-frequency noise in the force sensors

There is high-frequency noise in the force platform measurements with a standard deviation of 0.13 N.m for moments and 0.23 N for forces. We estimate the position of the CoM according to:

$$\frac{x_{CoM}}{L} = \frac{M_y}{L.M.g} + \frac{F_x}{M.g}$$

Across subjects, the smallest value of $M$ is 49.4 kg, and the smallest value of $M.L$ is 41.0 kg.m (corresponding to a weight of 49.4 kg and a limb length of 0.83 m). The standard deviation of this sum is thus lower than:

$$\sqrt{(\frac{0.13}{41.0 \times 9.8})^2 + (\frac{0.23}{49.4 \times 9.8})^2} = 0.00058$$

The instantaneous error in measurement is thus about 0.6 % limb length. This noise being however of high frequency, it is mostly abolished by taking averages over a 500 ms period (corresponding to 500 samples).

b) Low-frequency postural shifts

The main source of error in the estimation of $x_{CoM}$ comes from the subjects' small amplitude shifts in posture throughout the waiting period, which violates our assumption of stationarity. In CSRT trials, stationarity implies that the average value of $\frac{F_x}{M.g}$ is null, such that the mean position of $x_{CoM}$ is equal to the mean position of $x_{CoP}$. To evaluate the effect of non-stationarity, we therefore consider the offset between the measured values of initial $x_{CoM}$ and $x_{CoP}$ positions in CSRT trials, which corresponds to the mean value of $\frac{F_x}{M.g}$ in the 500 ms preceding trial onset. This has a mean value of 0.22 % limb length and a standard deviation of 0.18 % limb length (Figure 4). The bias of 0.22 % limb length suggests that the calibration of the force platforms is slightly inaccurate. Note that this bias is less than half of the instantaneous measurement noise.

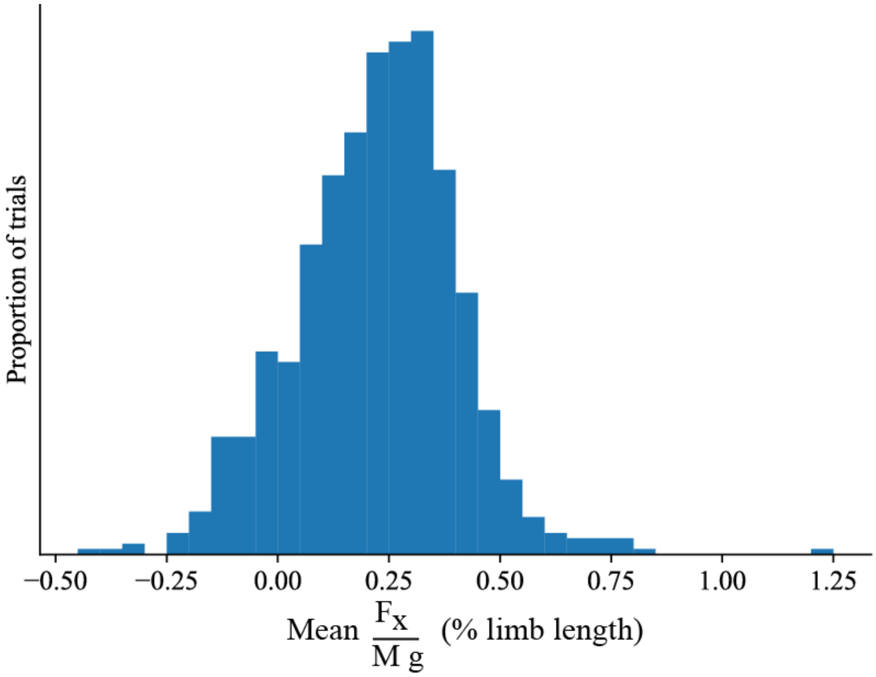


*Figure 4 Non-stationarity in CSRT trials in the 500 ms preceding trial onset*

c) Total error of measurement

The error of measurement of $x_{CoM}$ on a given trial has a constant offset due to the inaccurate calibration of the force platforms (which we estimate at around 0.22 % limb length), and errors in positioning the markers on the anatomical landmarks. It also has a variable noise component due to low frequency postural shifts (which we estimate at around 0.18% of limb length) and the error in measuring the position of the markers (which we estimate to be less than 1 mm, i.e. 0.1 % limb length). We thus estimate this variable noise component to be less than 0.3 % limb length.

3. Statistics

a) Proportion of trials without steps

The 2-proportion z-test is used to compare the proportion of non-step trials across subject groups and across perturbation amplitudes. The null hypothesis is that the observations in the two subsets come from two binary distributions with the same mean. With the first subset containing $N_{trials,1}$ trials and $N_{non\text{-}step,1}$ non-step trials, and the second subset containing $N_{trials,2}$ trials and $N_{non\text{-}step,2}$ non-step trials, the pooled sample mean is given by:

$$M = \frac{N_{nonstep,1} + N_{nonstep,2}}{N_{trials,1} + N_{trials,2}}$$

The standard error of the sample mean, when sampling $N_{trials}$ trials from a binary distribution of mean $M$ is given by:

$$SE = \sqrt{\frac{M(1-M)}{N_{trials}}}$$

Thus, the standard error of observing the two sample means is:

$$SE = \sqrt{\frac{M(1-M)}{N_{trials,1}} + \frac{M(1-M)}{N_{trials,2}}}$$

The test statistic is a z-score, which quantifies how likely the observed difference in proportions across the two subsets is, given this standard error:

$$z = \frac{\frac{N_{nonstep,1}}{N_{trials,1}} - \frac{N_{nonstep,2}}{N_{trials,2}}}{SE}$$

When the each subset of trials contains at least 10 successes and 10 failures (a condition which is indeed verified in our data), then this z-score is normally distributed, and the probability of observing a z-score at least as extreme is evaluated as a p-value.

For graphical representation (main text, Figure 2.E), we estimate the proportion of trials in a given subset as:

$$P = \frac{N_{nonstep,i}}{N_{trials,i}}$$

And the standard error on this estimate as:

$$SE = \sqrt{\frac{P(1-P)}{N_{trials,i}}}$$

b) Differences in medians of distributions

We use the Mann-Whitney U-test to evaluate whether there are differences in the medians of the distributions of:

- performance in perturbation trials with steps, between subject groups and between perturbation amplitudes
- initial CoM position between the CSRT task and the perturbation task, for each subject group
- initial CoM position between perturbation trials with and without steps, for each subject group.

The null hypothesis is that the observations come from distributions with the same median. To evaluate this, the observations are pooled and ranked, and $R_1$ the sum of ranks of the first group is calculated . The U-statistic is given by:

$$U = R_1 - \frac{N_{trials,1}(N_{trials,1} + 1)}{2}$$

Note that if the samples from the first group are all smaller than those from the second group, then $R_1 = \frac{N_{trials,1}(N_{trials,1}+1)}{2}$, and $U = 0$.

The ranking of observations is illustrated in Figure 5.A for the comparison of performance in perturbation trials with steps, between elderly (white) and young (black) subjects, for short (left panel) and long (right panel) perturbations.

When a statistical difference in medians is found, then the amplitude of this difference is estimated using the Hodges-Lehmann estimator. All possible pairs are formed with a sample from the first and a sample from the second distribution. For each pair, the difference between the first and second sample is calculated, and the median of these differences is used as an estimate of the difference between the two distributions. The distribution of differences is illustrated in Figure 5.B for the comparison of performance in perturbation trials with steps, between elderly and young subjects, for short (left panel) and long (right panel) perturbations.

When no statistical difference in medians is found, then the Kolmogorov-Smirnov test is used to determine whether there is any difference in the distributions. The null hypothesis is that the two sets of observations come from an identical distribution. The test statistic is the maximal difference between the cumulative distribution functions of the two observations, shown as a black bar in Figure 5.C for the comparison of performance in perturbation trials with steps, between elderly non-fallers (blue) and elderly fallers (red), for short (left panel) and long (right panel) perturbations.

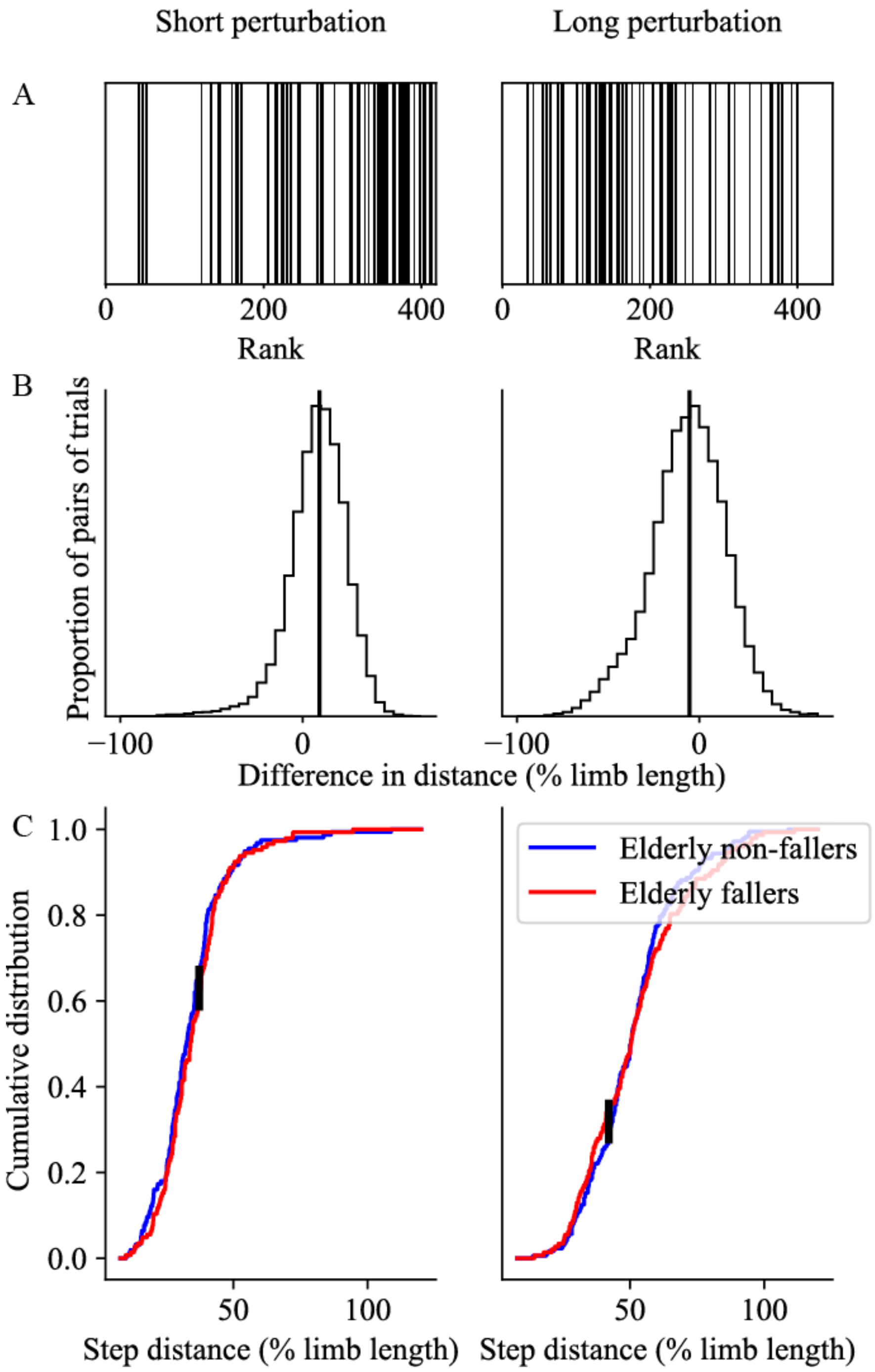


*Figure 5 Illustration of statistical methods. Differences in the distribution of distances required to stop in perturbation trials with steps (normalised to limb length), between young and elderly subjects (A,B), and between elderly fallers (red) and elderly non-fallers (blue) (C), for short perturbations (left panels) and long perturbations (right panels). A. Trials sorted by rank, with the elderly shown in white and the young shown in black. B. Distribution of the difference in distance between elderly and young subjects, for all pairs of trials. C. Cumulative distribution of distances for elderly non-fallers (blue) and elderly fallers (red). The maximal difference between the two cumulative distributions is indicated as a black bar.*
